# Altered visual function in a larval zebrafish knockout of neurodevelopmental risk gene *pdzk1*

**DOI:** 10.1101/2020.09.21.307405

**Authors:** Jiaheng Xie, Patricia R. Jusuf, Bang V. Bui, Stefanie Dudczig, Patrick T. Goodbourn

**Affiliations:** School of BioSciences, The University of Melbourne; Department of Optometry and Vision Sciences, The University of Melbourne; Melbourne School of Psychological Sciences, The University of Melbourne

**Keywords:** PDZK1, visual phenotyping, zebrafish, optomotor response (OMR), electroretinogram (ERG), neurodevelopmental disorders

## Abstract

The human *PDZK1* gene is located in a genomic susceptibility region for neurodevelopmental disorders. A genome-wide association study (GWAS) identified links between *PDZK1* polymorphisms and altered visual contrast sensitivity, an endophenotype for schizophrenia and autism spectrum disorder. The PDZK1 protein is implicated in neurological functioning, interacting with synaptic molecules including post-synaptic density 95 (PSD-95), *N*-methyl-D-aspartate receptors (NMDAR), corticotropin-releasing factor receptor 1 (CRFR1) and serotonin 2A receptors. To elucidate the role of PDZK1, we generated *pdzk1-*knockout (*pdzk1-*KO) zebrafish using CRISPR/Cas-9 genome editing. Visual function of 7-day-old fish was assessed at behavioural and functional levels using the optomotor response (OMR) and scotopic electroretinogram (ERG). We also quantified retinal morphology and densities of PSD-95, NMDAR1, CRFR1 and serotonin in the synaptic inner plexiform layer at 7 days, 4 weeks and 8 weeks of age. Relative to wild-type, *pdzk1*-KO larvae showed spatial-frequency tuning functions with increased amplitude (likely due to abnormal gain control) and reduced ERG *b*-waves (suggestive of inner retinal dysfunction). However, these functional differences were not associated with gross synaptic or morphological retinal phenotypes. The findings corroborate a role for *pdzk1* in visual function, and our model system provides a platform for investigating other genes associated with abnormal visual behaviour.

## 1. Introduction

Altered visual contrast sensitivity is a visual endophenotype of many neurodevelopmental disorders, including schizophrenia and autism spectrum disorder (ASD) [1, 2]. A genome-wide association study (GWAS) identified a link between abnormal contrast sensitivity and the single-nucleotide polymorphism (SNP) rs1797052, located in the 5′-untranslated region of the *PDZK1* gene, which is situated in a high-risk locus for schizophrenia and ASD [3]. Variation in this gene may thus contribute to visual deficits observed in a number of different disorders.

As a scaffolding protein, PDZK1 is implicated in neurosynaptic signalling. PDZK1 contains four tandem post-synaptic density 95/disk-large/ZO-1 (PDZ) domains [4], which allows it to interact with proteins possessing PDZ-binding regions. For example, PDZK1 binds with post-synaptic density 95 (PSD-95), *N*-methyl-D-aspartate receptors (NMDAR), Synaptic Ras GTPase-activating protein 1 (SynGAP1) and Kelch-like protein 17 (KLHL17) to form a complex that anchors proteins to the cytoskeleton on cell membranes [5]. Like PDZK1, the PSD-95 protein also contains PDZ domains, and functions as a scaffold protein on post-synaptic membranes for excitatory synapses [6], while KLHL17 is a brain-specific Kelch protein that binds to intracellular F-actin, a major component of the cellular cytoskeleton [7]. PDZK1 thus likely plays a role in maintaining the distribution and shape of post-synaptic densities by regulating actin-based neuronal functions [7]. Disruption of PSD-95 (also called DLG4) also affects the Neurexin-Neuroligin-Shank (NRXN-NLGN-SHANK) pathway, leading to cognitive dysfunction [8, 9].

PDZK1 is important for anchoring and clustering NMDARs, ionotropic glutamate receptors responsible for gating ion influx into neurons [10]. In the visual system, NMDARs regulate contrast gain control, optimising visual perception of contrast and motion, and abnormalities in these receptors have been hypothesised to underpin visual disturbances in schizophrenia [10, 11]. PDZK1 also regulates signalling and endocytosis for the corticotropin-releasing factor receptor 1 (CRFR1) and the serotonin 2A receptor (5-HT2AR) [12]. CRFR1 is a vital component of the hypothalamic-pituitary-adrenal axis for stress responses [13], and it sensitises serotonin 2 receptors including 5-HT2AR to modulate anxiety-related behaviours [14]. PDZK1 thus interacts with multiple membrane-associated proteins for synaptic signalling, with likely impacts on neural processing, visual perception and behaviour. However, despite evidence of genetic associations, to date there has been no direct demonstration of an impact of *PDZK1* disruption on behaviour and physiology.

The retina at the back of the eye is an extension of the central nervous system (CNS) that provides a highly accessible readout of gene function in the CNS. Neurodegeneration and neurodevelopmental abnormalities in the brain and spinal cord typically manifest in visual dysfunction, and some CNS disorders (e.g. stroke, multiple sclerosis, and Alzheimer disease) can be diagnosed by ocular symptoms [15]. Compared to the brain, the retina has a relatively straightforward organisation with well characterised excitatory (photoreceptors, bipolar and ganglion cells) and inhibitory (horizontal and amacrine cells) neuronal classes arranged in distinct organised layers. The visual system is highly amenable to assessment, with robust non-invasive behavioural (e.g. *optomotor response*; OMR), physiological (e.g. *electroretinogram*; ERG) and histological tools that allow the neurological function of genes to be comprehensively evaluated across these different phenotypic levels.

The zebrafish (*Danio rerio*) has key advantages as a model for studying both visual and gene function, including conserved retinal architecture and genetics with other vertebrates, ease of genetic manipulation, high fecundity and rapid development. Importantly, as in humans, a single *pdzk1* gene is expressed in larval zebrafish retina and brain [16]. Here, we characterised the effects of *pdzk1* disruption using CRISPR/Cas-9 genome modification to generate a *pdzk1* knockout (*pdzk1*-KO) zebrafish mutant. Visual behaviour was assessed by quantifying spatial-frequency tuning and contrast-response functions using OMR, and physiological function of the retina was examined with ERG. To investigate possible histological correlates of the observed behavioural and functional phenotypes, we quantified general retinal morphology via retinal size, and thickness of the whole retina or synaptic inner plexiform layer (IPL), as well as the density of PSD-95, NMDAR1, CRFR1 and serotonin in the IPL. We found that relative to wild-type larvae, *pdzk1*-KO larvae showed abnormal contrast gain control, with significantly lower contrast thresholds that lead to an increase in the overall amplitude of the spatial-frequency tuning function. In addition, *pdzk1*-KO larvae showed clear deficits in retinal function, with a reduction in the ERG *b*-wave amplitude suggesting dysfunction of the retinal bipolar cells. We found no clear differences between *pdzk1*-KO and wild-type fish in anatomical and histological markers at 7 days post-fertilisation (dpf), 4 weeks post-fertilisation (wpf) and 8 wpf, ruling out a number of the most likely candidates for the anatomical substrate of *pdzk1*’s effects on visual function.

## 2. Methods

### 2.1 Animal husbandry

Zebrafish *(Danio rerio*) were maintained and bred in the Fish Facility at the Walter and Eliza Hall Institute of Medical Research according to local animal guidelines. For experiments at 7 dpf, embryos and larvae (prior to sex determination) were grown in Petri dishes in an incubator at 28.5°C before use. Fish for experiments at higher ages were grown in Petri dishes at 28.5°C up to 5 dpf and then introduced to tanks and raised in flow-through systems at 28°C. All procedures were performed according to the provisions of the National Health and Medical Research Council *Australian Code of Practice for the Care and Use of Animals for Scientific Purposes* (2013) [17]. They were approved by the Faculty of Science Ethics Committee at the University of Melbourne (application number 1614054.4).

### 2.2. Generation of mutants

To disable *pdzk1* gene function, we applied a *clustered regularly interspaced short palindromic repeats* (CRISPR)/CRISPR-associated (Cas) system according to published protocols [18]. The guide RNA (GAAGGTAGAAGCCATAACCC followed by a short hairpin RNA scaffold), Cas-9 enzyme (Genesearch, Arundel, QLD, Australia) and stop-codon cassette (GTCATGGCGTTTAAACCTTAATTAAGCTGTTGTAG) with homologous sequence hands were co-injected into zebrafish zygotes. Homology-directed repair (HDR) resulted in a premature stop codon at exon 2 of the *pdzk1* gene, leading to the loss of functional PDZK1 in mutants (Supplementary Figure S1a–a″). To highlight cell layers in the zebrafish retina for quantification of retinal morphology, the Tg(*ptf1a*:*GFP*) line (provided by Steven D. Leach, John Hopkins University, Baltimore, MD) was crossed with the *pdzk1*-KO mutants to generate a *pdzk1*-KO/Tg(*ptf1a*:*GFP*) line. The transgenic construct labels with green fluorescent protein the horizontal and amacrine cells [19, 20], which project into outer and inner plexiform layers, respectively.

### 2.3 Genotyping

For DNA isolation, small wedges of fin were cut from anaesthetised 3-month-old fish. Fin samples were incubated in lysis buffer (10 mM Tris HCl, 50 mM KCl, 0.3% Tween and 0.3% IGEPAL; pH 8.3) with 1.25 mg/ml Proteinase K (Roche, Bella Vista, NSW, Australia; cat. number 03115879001) at 55°C for 2.5 h, followed by incubation at 98°C for 10 min. Isolated DNA was used as a template for polymerase chain reaction (PCR) amplification of the targeted *pdzk1* genomic DNA sequence (*Genomic DNA primers*; Supplementary Table S1). The PCR program comprised an initial denaturation step of 98°C for 3 min, 35 cycles of 98°C for 30 s, 60°C for 30 s and 72°C for 1 min, followed by a final extension step of 72°C for 3 min. Electrophoresis of the PCR products was performed using 3% Tris-acetate-EDTA (TAE) agarose gel. Target bands were cut from electrophoresis gels and the PCR products were extracted using a QIAquick gel extraction kit (QIAGEN, Hilden, Germany). Extracted products were amplified by PCR using a forward primer (*Primer for sequencing*; Supplementary Table S1) and sent to Macrogen (Seoul, South Korea) for sequencing (Supplementary Figure S1b–b′). The mutants used in this study were all homozygous *pdzk1*-KO zebrafish, including those expressing the Tg(*ptf1a*:*GFP*) construct.

### 2.4 Optomotor response (OMR)

#### (a) Apparatus and procedure

The OMR apparatus was adapted from one previously described (Supplementary Figure S2) [21, 22]. Visual stimuli were generated by a Power Mac G5 computer (Apple Inc., Cupertino, CA, USA) running MATLAB R2016b (MathWorks, Natick, MA, USA) with Psychtoolbox extensions [23] and were processed on an ATI Radeon HD 5770 graphics card (AMD, Santa Clara, CA, USA). The outputs were sent to a BITS++ video processor (Cambridge Research Systems, Rochester, UK) for increased contrast resolution, and displayed on a cathode ray tube (CRT) monitor (Model M992, Dell Inc., Round Rock, TX, USA) with its screen facing upwards. 7-dpf larvae (*N* = 50–58 in each group) were contained in a six-lane arena with a transparent base (Supplementary Figure S2b), positioned above the monitor screen. Test stimuli were displayed on the screen below the arena, drifting at 25, 50 or 100 degrees per second (°/s) parallel to the long axis of the lanes during a trial (Supplementary Figure S2a). Before experiments, larvae were transferred to arena lanes within 10 min, after which they were allowed to adapt to the arena for 10 min. Prior to each trial, a corralling stimulus (25 °/s drift) was shown for 30 s to guide larvae to the centre of the lane (Supplementary Figure S2d). This was followed by a 30-s presentation of a test stimulus. A C922 Pro Stream webcam (Logitech Company, Lausanne, Switzerland) fixed over the arena was controlled by a custom MATLAB program to take digital images before and after each presentation of test stimuli for calculation of swimming distance. A blank grey screen was presented after the offset of the test stimulus as the texture for the next trial was computed.

For measurement of spatial-frequency tuning functions, test stimuli were Gaussian noise textures filtered to be spatial-frequency bandpass with centre frequencies of 0.005, 0.01, 0.02, 0.04, 0.08, 0.16 or 0.32 cycles per degree (c/°; *SD* = 0.5 octaves) at full contrast (100%). There were 72 trials for each genotype group (*pdzk1-KO* and wild-type), per combination of spatial frequency and speed. For measurement of contrast-response functions, the centre spatial frequency was 0.02 c/° (*SD* = 0.5 octaves), with textures presented at 1%, 3%, 5%, 10%, 30%, 50%, or 100% contrast. Wild-type and mutant groups performed 36 and 24 trials, respectively, per combination of contrast and speed. All experiments were conducted between 9:30 a.m. and 6:00 p.m. After experiments, larvae were humanely killed using 0.1% tricaine (Sigma-Aldrich, St Louis, MA, USA, cat. number E10521-50G).

#### (b) OMR image analysis

Larval positions were extracted from before and after images for each trial using a custom MATLAB algorithm. Image contrast was flattened, and a Laplacian-of-Gaussian filter was applied to highlight edges [24]. Larvae were segmented by thresholding (manually adjusted if necessary), and the centroid of larval positions was calculated using *bwconncomp* and *regionprops* commands from the MATLAB Image Processing Toolbox. On each trial, the change in position of the centroid in the direction of the texture motion was computed as the optomotor index (OMI).

Normalised OMIs for spatial-frequency tuning functions were calculated by normalising all data to the OMI of wild-type larvae for 100%-contrast, 0.02-c/° stimuli drifting at 25 °/s (i.e., the condition typically producing the greatest response). The spatial-frequency tuning function was fitted as a log-Gaussian using a least-squares criterion. For each fitted model, the estimated parameters were amplitude (i.e., height of the peak), peak spatial frequency (i.e., spatial frequency at which amplitude peaked) and bandwidth (i.e., the standard deviation). Normalised OMIs for contrast-response functions were calculated by normalising all data to the OMI of the wild-type group for 100% contrast at 50 °/s and 0.02 c/°. The contrast-response function was fitted as a two-parameter piecewise function (Equation 1) using a least-squares criterion:

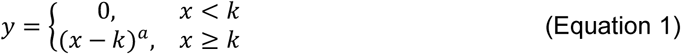

where *y* is the OMI, *x* is log stimulus contrast, *k* is contrast threshold (i.e., the minimum contrast to evoke an optomotor response) and *a* is response gain (i.e., the slope of the function).

To test whether spatial-frequency tuning and contrast-response functions differed between groups, an omnibus *F*-test was used to compare a full model, in which parameter estimates of each group could vary independently, with a restricted model, in which parameters were constrained to be the same across groups. To determine whether specific parameter estimates differed between groups, a nested *F*-test was used to compare a full model with a restricted model in which one parameter was constrained to be the same across groups [25]. A criterion of α = 0.05 was used to determine significance, with Bonferroni correction applied to *P* values to account for multiple testing where appropriate.

### 2.5 Electroretinography (ERG)

Scotopic ERGs were measured in 7-dpf fish according to our published method [26]. Larvae were dark-adapted (>8 h, overnight) prior to experiments and anaesthetised using 0.02% tricaine in 1× goldfish ringer’s buffer. For testing, an individual larva was transferred onto a moist PVA sponge platform. Under dim red illumination (17.4 cd.m^-2^, λ_max_ 600 nm), a sponge-tipped recording electrode (<40 µm diameter) gently touched the central corneal surface of the larval eye and the reference electrode was inserted into the sponge platform. After electrode placement, the platform was inserted into a Ganzfeld bowl and the larva was allowed to dark adapt for >3 min. ERG responses were recorded from 22 wild-type and 26 *pdzk1*-KO larvae with flash stimuli at −2.11, −0.81, 0.72, 1.89 or 2.48 log cd.s.m^−2^. At −2.11 and −0.81 log cd.s.m^−2^, three repeats were measured with an inter-flash interval of 10 s. At 0.72 to 2.48 log cd.s.m^−2^, a single response was measured with 60 s between flashes. All experiments were performed between 9:00 am and 6:00 pm at room temperature. Larvae were humanely killed after experiments using 0.1% tricaine. Amplitudes of the *a*- and *b*-waves were measured from baseline to the negative *a*-wave trough and from the negative *a*-wave trough to the *b*-wave peak, respectively. Implicit times of the *a*- and *b*-waves were measured from stimulus onset to the *a*-wave trough and the *b*-wave peak, respectively. Two-way ANOVA with Bonferroni correction was performed in Prism 7 (GraphPad, San Diego, CA, USA; α = 0.05). Best-fit lines were derived from a four-parameter logistic (4PL) sigmoidal function.

### 2.6 Histology

#### (a) Immunohistochemistry

Whole zebrafish at 7 dpf or 4 wpf, or dissected zebrafish eyes at 8 wpf, were fixed in 4% paraformaldehyde (PFA) in phosphate-buffered saline (PBS) for 3 h at room temperature, or overnight at 4°C. They were cryoprotected in 30% sucrose in PBS, embedded in OCT (Tissue-Tek) and cryo-sectioned (12 µm; Leica CM 1860 Cryostat, Wetzlar, Germany). For wild-type and *pdzk1*-KO larvae, antibody staining was carried out at room temperature using standard protocols. Antigen retrieval was performed by incubating slides in boiled 10 mM sodium citrate (pH 6) until the solution was cooled to room temperature. Slides were blocked in 5% foetal bovine serum (FBS) for 30 min and incubated overnight in rabbit anti-PSD-95 (1:100; Abcam, Cambridge, United Kingdom; cat. number ab18258), anti-CRFR1 (1:100; Abcam; cat. number ab59023), anti-GluN1 (1:400; Synaptic systems; cat. number 114 011) or anti-serotonin (1:500; Abcam; cat. number ab66047) primary antibodies diluted in FBS. Slides were subsequently incubated for 2 h in secondary antibodies (all 1:500; Thermo Fisher Scientific, Mulgrave, Victoria, Australia) diluted in 5% FBS. The secondary antibodies used were anti-rabbit Alexa Fluor-488 (cat. number A11008) or Alexa Fluor-546 (cat. number A11010), anti-mouse Alexa Fluor-488 (cat. number A11001) and anti-goat Alexa Fluor-488 (cat. number A11055). Nuclei were counterstained with 4′,6-diamidino-2-phenylindole (DAPI; 1:10000; Sigma-Aldrich, St. Louis, MO, USA; cat. number D9542-10MG) in PBS for 20 min and sections were mounted in Mowiol (Sigma-Aldrich; cat. number 81381-250G). For Tg(*ptf1a*:*GFP*) and *pdzk1*-KO/Tg(*ptf1a*:*GFP*) larvae, samples were stained with 4′,6-diamidino-2-phenylindole (DAPI) only, following cryostat sectioning.

#### (b) Retinal image acquisition

For quantification of IPL densities, retinae stained with antibodies for PSD-95, NMDAR1, CRFR1 and serotonin from *pdzk1*-KO and wild-type larvae were imaged using a Nikon A1R confocal microscope (Nikon, Tokyo, Japan) with a 60× or 40× oil objective lens. For quantification of IPL thickness, retinal size and retinal thickness, images of retinal sections of Tg(*ptf1a*:*GFP*) and *pdzk1*-KO/Tg(*ptf1a*:*GFP*) larvae were taken within two sections from the optic nerve, using a ZEISS Axioscope (Carl Zeiss Microscopy GmbH, Oberkochen, Germany) with a 20× objective lens, or a Nikon A1R confocal microscope with a 40× oil objective lens. For all confocal imaging, the *deconvolution* function was applied to minimise background noise.

#### (c) Retinal image analysis

To measure puncta density, three regions of interest (ROI; 10 × 20 µm) were randomly selected from the IPL of one confocal image for each retina using FIJI [27]. For each ROI, thresholding was performed with the *invert LUT* function to select areas containing no puncta and create a background mask. All potential puncta in the ROI were identified as dots using the *Find Maxima* function. Dots identified in the background area (i.e., noise) were masked using the *Image Calculator* function and the background mask. The remaining dots in the ROI were considered true puncta and quantified for analysis. The density of the analysed molecule in each retinal image was quantified as the average of three ROIs. Statistical analysis for puncta density was performed using two-way ANOVA with Bonferroni correction (Prism 7; α = 0.05). To quantify evidence for null hypotheses, we also calculated Bayesian ANOVA using JASP (*BF*_inclusion_ = 0.33; i.e., the data are at least three times as likely under the null hypothesis than the alternative) [28]. For retinal size, the *polygon selection* function was used to outline the whole retinal section of Tg(*ptf1a*:*GFP*) and *pdzk1*-KO/Tg(*ptf1a*:*GFP*) images, and the size of the outlined area was measured using FIJI [27]. IPL thickness was measured by a masked observer in three locations (central and 45° on either side) using FIJI. For each eye, the average over the three locations was taken as the IPL thickness. Retinal thickness was similarly measured as the average retinal thickness in the three locations. Retinal size, IPL thickness and retinal thickness were normalised to the mean of wild-type group parameters. Morphological distributions were approximately normal and were analysed using unpaired *t-*tests (Prism 7; α = 0.05). There were 12–31 retinae per group for puncta density analysis (Supplementary Table S2), and 12 (wild-type) or 13 (*pdzk1*-KO) retinae for morphological analysis.

## 3. Results

### 3.1 Spatial-frequency tuning functions

To determine the contribution of *pdzk1* to spatial vision, we measured the spatial-frequency tuning function using OMR in wild-type and *pdzk1*-KO larvae generated using CRISPR/Cas-9 genome modification (Supplementary Figure S1). All spatial-frequency tuning functions were well fit by a log-Gaussian model (Figure 1a–c). In an omnibus test (Supplementary Table S3), overall spatial-frequency tuning functions statistically differed between wild-type and *pdzk1*-KO larvae at each speed (Figure 1a– c). Examining individual parameters, there was a significantly higher amplitude (i.e., height of the spatial-frequency tuning function peak; Figure 1d; Supplementary Table S3) for mutants compared with wild-type fish at each speed tested. No differences were found in the tuning function’s peak frequency (Figure 1e) or bandwidth (standard deviation; Figure 1f) at any speed (Supplementary Table S3).

**Figure 1.**
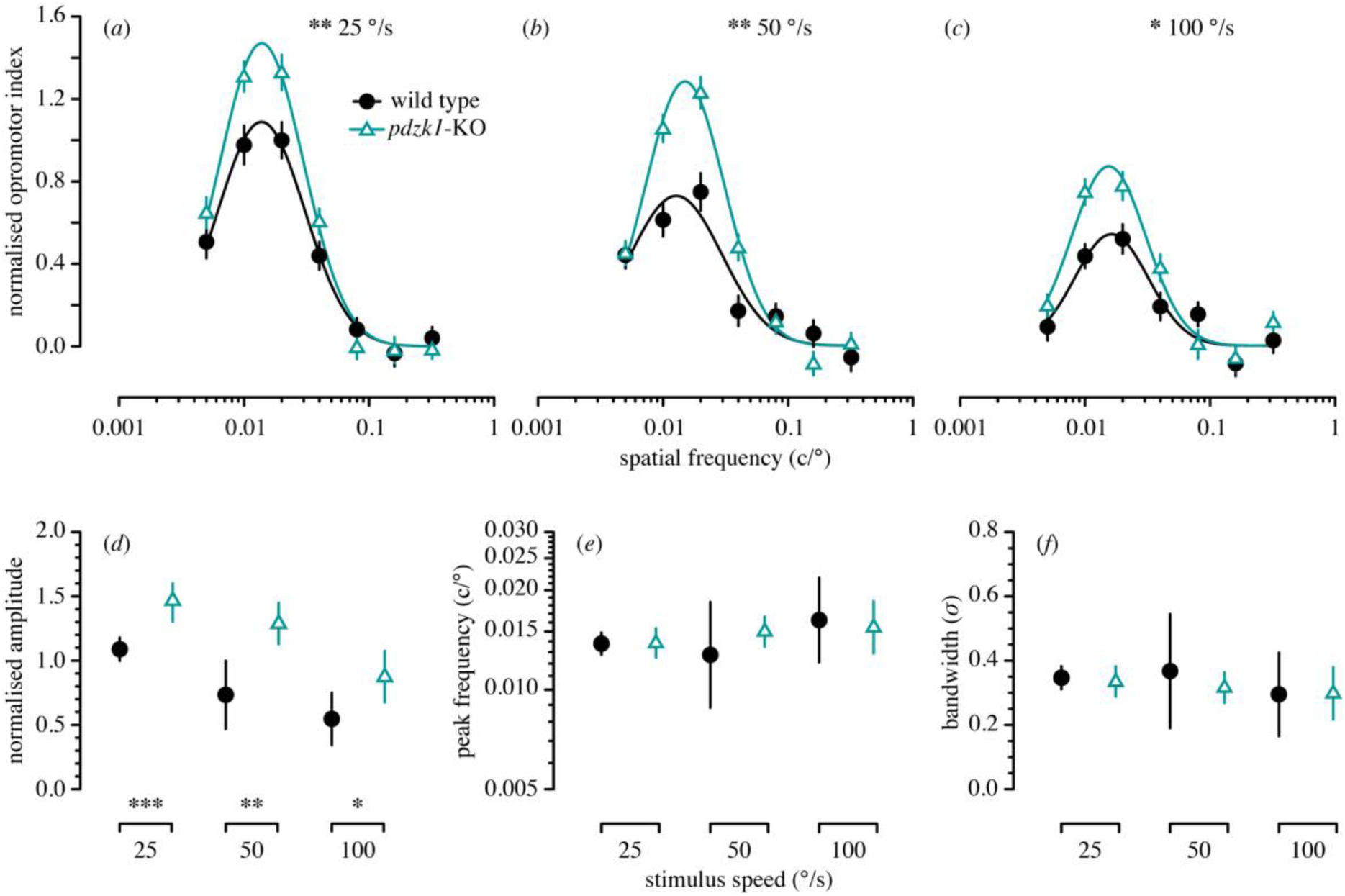
Spatial-frequency tuning functions measured using the optomotor response (OMR) of wild-type and *pdzk1* knockout (*pdzk1*-KO) larvae. The upper panels show normalised optomotor index as a function of spatial frequency for wild-type (black circles and lines) and *pdzk1*-KO larvae (cyan triangles and lines) at (**a**) 25, (**b**) 50 and (**c**) 100 °/s. Spatial-frequency tuning functions are three-parameter log-Gaussian functions fit to the data by minimising the least-squares error. Error bars show ±SEM across trials. The bottom panels show (**d**) normalised amplitude, (**e**) peak spatial frequency and (**f**) bandwidth as a function of group at the three stimulus speeds tested. Error bars show 95% confidence intervals on the fitted parameter. \**P* < 0.05; \*\**P* < 0.01; \*\*\**P* < 0.001.

### 3.2 Contrast-response functions

To examine whether the increased amplitude of the spatial-frequency tuning function in *pdzk1*-KO mutants was due to altered contrast gain (i.e., decreased threshold) or response gain (i.e., increased motor response to supra-threshold stimuli), we used OMR to measure the contrast-response function at 0.02 c/°, at which zebrafish larvae typically showed the most robust responses. Contrast-response functions were well fit by a two-parameter piecewise function (Figure 2). In omnibus analysis, functions were significantly different between groups at 25, 50 and 100 °/s (*P* < 0.001, *P =* 0.006 and *P* = 0.001), respectively; Figure 2a–c; Supplementary Table S4). Examining individual parameters, we found no difference in response gain (function slope) between wild-type and mutant groups at any tested speed (Figure 2d; Supplementary Table S4). However, contrast threshold was significantly lower for mutants than for wild-type larvae at 25 and 50 °/s (*P* < 0.001 and *P* = 0.002, respectively; Figure 2e and Table S4), indicating a higher contrast sensitivity for mutants. Overall, all OMR contrast-response functions deviated upwards at lower contrasts for mutants than for wild-type larvae, suggesting that the augmented spatial-frequency tuning functions in mutants reflect increased contrast gain rather than response gain.

**Figure 2.**
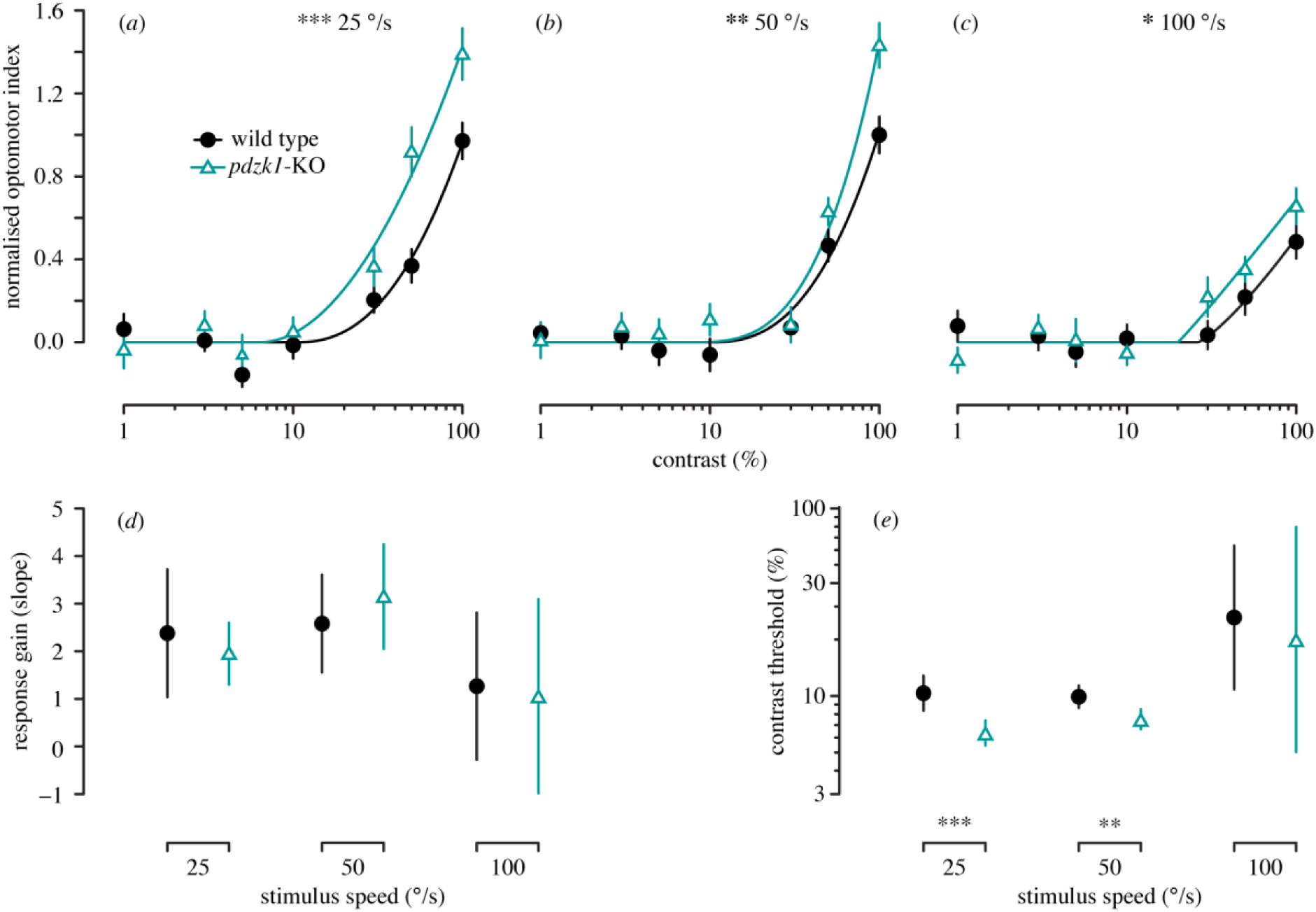
Contrast-response functions measured using the optomotor response (OMR) in wild-type (black circles and lines) and *pdzk1* knockout (*pdzk1*-KO) larvae (cyan triangles and lines) at (**a**) 25, (**b**) 50 and (**c**) 100 °/s. Contrast-response curves are fit to the data using two-parameter piecewise functions by minimising the least-squares error with respect to the data. Error bars show ± SEM across trials. The bottom panels show (**d**) response gain and (**e**) contrast threshold as a function of group at the three stimulus speeds tested. Error bars show 95% confidence intervals on the fitted parameter. \**P* < 0.05; \*\**P* < 0.01; \*\*\**P* < 0.001.

### 3.3 Electroretinography (ERG)

We conducted scotopic ERG measurements for wild-type controls and mutant larvae. Compared to wild-type, *pdzk1*-KO showed significantly smaller *b*-wave amplitudes (*P* < 0.0001), particularly at higher stimulus intensities (*P* = 0.032 and *P* < 0.0001 for 1.89 and 2.48 log cd.s.m^−2^, respectively; Figure 3a and 3d; Supplementary Table S5). There was no statistical difference in *a*-wave amplitudes, nor in implicit times of *a*- and *b*-waves, between wild-type and *pdzk1*-KO mutants (Figure 3a–c and 3e; Supplementary Table S5).

**Figure 3.**
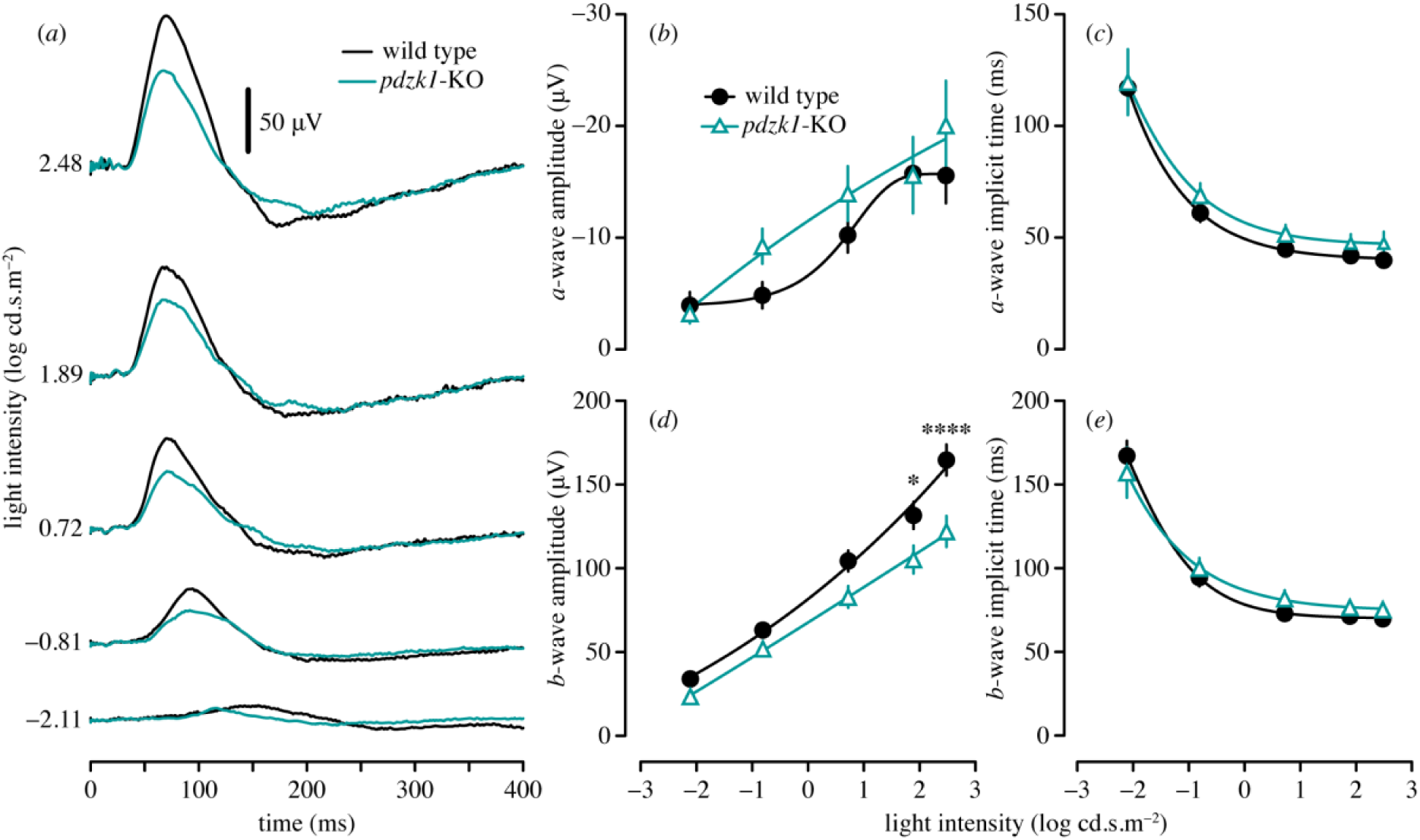
Scotopic electroretinograms (ERG) of wild-type and *pdzk1* knockout (*pdzk1*-KO) larvae. (**a**) Group average ERG traces. Wild-type and *pdzk1*-KO larvae responses are shown as black and cyan lines, respectively, at −2.11, −0.81, 0.72, 1.89 and 2.48 log cd.s.m^−2^. Scale bar shows 50 µV. Remaining panels show group average (± SEM) (**b**) a-wave amplitude, (**c**) a-wave implicit time, (**d**) b-wave amplitude and (**e**) b-wave implicit time for wild-type (black circles) and *pdzk1*-KO (cyan triangles) larvae. Lines were fit using a four-parameter sigmoidal function. Data were compared using two-way ANOVA with Bonferroni correction. \**P* < .05; \*\*\*\**P* < .0001.

### 3.4 Retinal morphology

Differences in retinal function might arise from changes to retinal morphology. To identify distinct layers in the zebrafish retina, we used the transgenic construct Tg(*ptf1a*:*GFP*) to genetically label horizontal and amacrine cells, and their projections into the OPL and IPL, respectively (Figure 4a–b″). Our quantification revealed no difference in IPL thickness, retinal size or retinal thickness in *pdzk1*-KO compared with wild-type larvae (Figure 4c–d; Supplementary Table S6).

**Figure 4.**
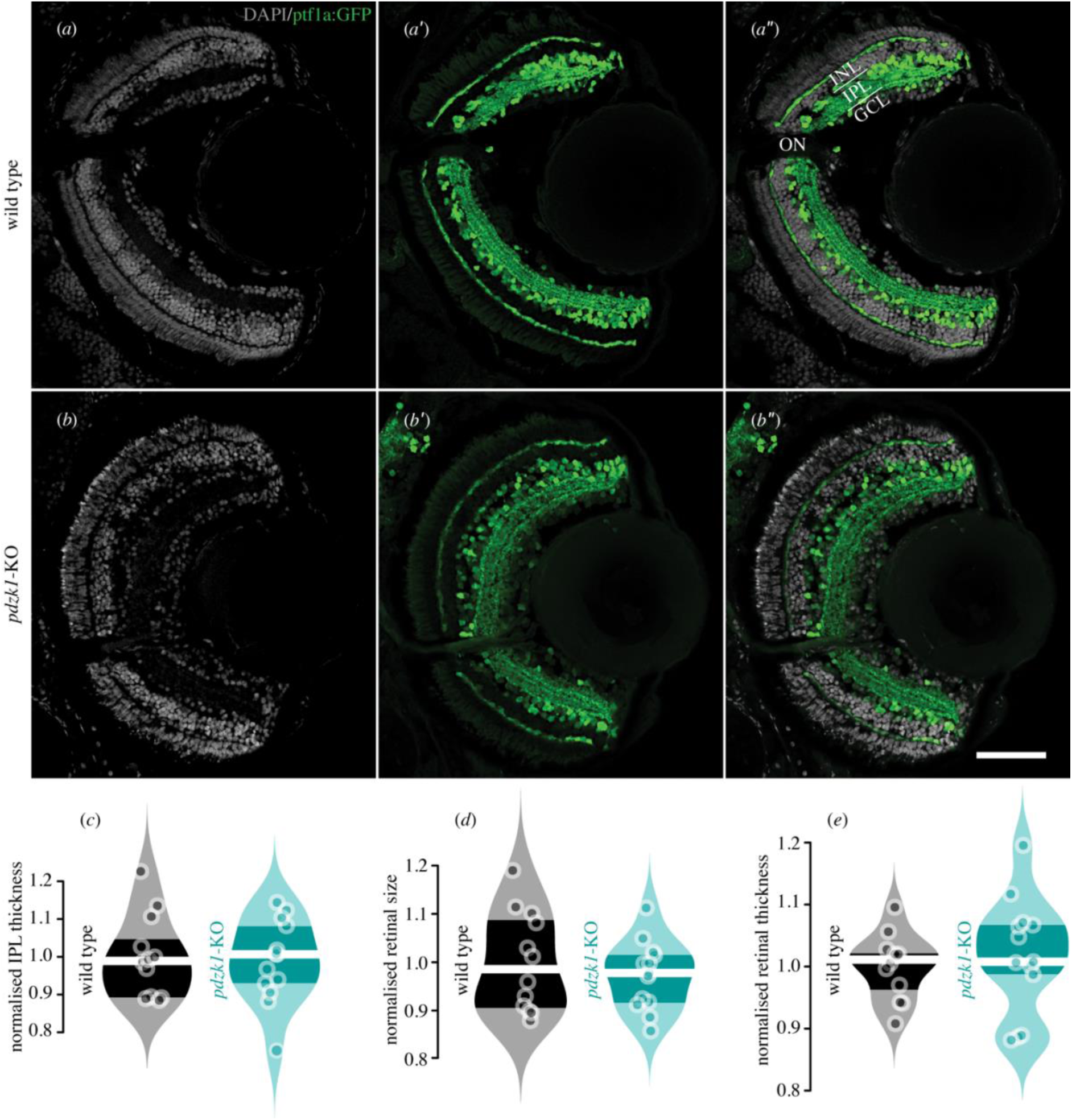
Transgenic labelling of retinal layers and quantification of retinal morphology for wild-type and *pdzk1* knockout (*pdzk1*-KO) larvae. (**a**–**b**″) Inhibitory neurons, including horizontal and amacrine cells, and their projections into the outer and inner plexiform layers were labelled by transgenic construct Tg(*ptf1a*:*GFP*) in green, revealing distinct retinal layers in wild-type and *pdzk1*-KO retinae. Nuclei stained with DAPI are shown in grey. Scale bar shows 50 µm. INL: inner nuclear layer; IPL: inner plexiform layer; GCL: ganglion cell layer; ON: optic nerve. Normalised (**c**) inner plexiform layer (IPL) thickness, (**d**) retinal size and (**e**) retinal thickness were quantified. Points are data from individual retinae. White lines represent medians and dark bands indicate inter-quartile ranges. For all analyses, unpaired *t*-tests were performed with 12 and 13 retinae for wild-type and *pdzk1*-KO groups, respectively.

### 3.5 PSD-95, serotonin, NMDAR1 and CRFR1 density in the retinal IPL

To investigate potential subcellular mechanisms underlying the observed changes in visual function, we quantified the density of synaptic signalling molecules known to interact with PDZK1 (i.e., PSD-95, serotonin, NMDAR1 and CRFR1) in the IPL at 7 dpf, 4 wpf and 8 wpf. As there is no commercially available antibody for the serotonin 2A receptor, we labelled serotonin to assess the effect of *pdzk1* knockout on the serotonin system. Interestingly, in both wild-type and *pdzk1*-KO larvae, our serotonin antibody rarely labelled serotonin in cell bodies, but there was strong serotonin co-labelling with PSD-95 on cell membranes and in synapses (Figure 5a–b″; Supplementary Figure S3a–d″); this serotonin antibody thus appears to mark serotonin receptors.

**Figure 5.**
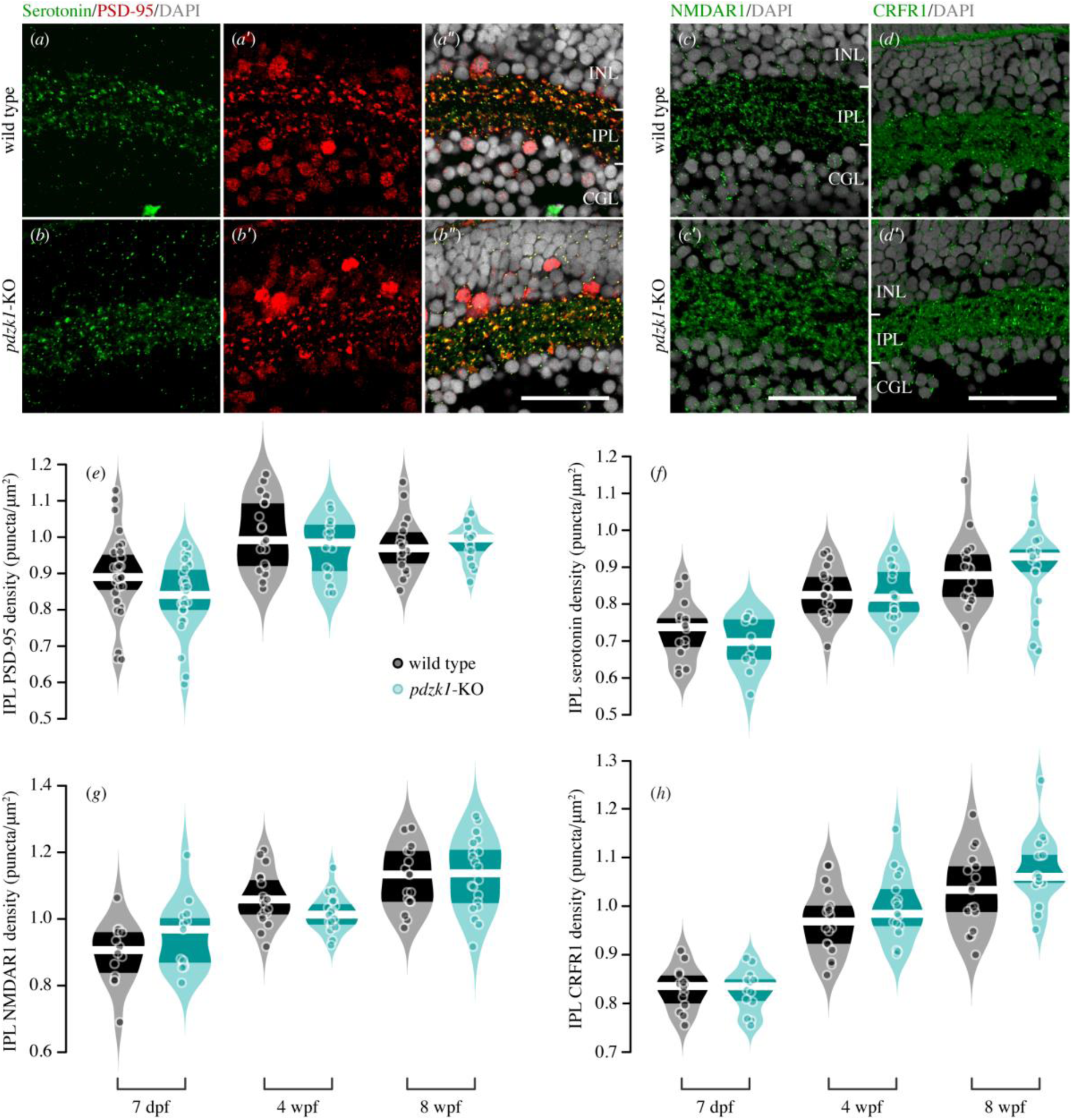
Immunostaining and density quantification of PDZK1-interacting molecules in wild-type and *pdzk1*-knockout (*pdzk1*-KO) retinae. Micrographs of retinal sections from 7 days post-fertilisation (dpf) wild-type and *pdzk1*-KO larvae were co-labelled with (**a, b″**) serotonin and post-synaptic density 95 (PSD-95) in green and red, respectively. (**c, c′**) *N*-methyl-D-aspartate receptor 1 (NMDAR1) and (**d, d**′) corticotropin-releasing factor receptor 1 (CRFR1) were labelled in green for 7 dpf retinae. For all images, nuclei were stained with DAPI, shown in grey. Scale bars represent 25 µm. PSD-95: post-synaptic density 95; NMDAR1: N-methyl-D-aspartate receptor 1; CRFR1: corticotropin-releasing factor receptor 1; INL: inner nuclear layer; IPL: inner plexiform layer; GCL: ganglion cell layer. Violin plots show densities of (**e**) PSD-95, (**f**) serotonin, (**g**) NMDAR1 and (**h**) CRFR1 in the IPL at 7 dpf, 4 wpf and 8 wpf. Points are data from individual retinae. White lines represent medians and dark bands indicate inter-quartile ranges. Statistical comparisons were performed using two-way ANOVA with Bonferroni correction and Bayesian ANOVA. There were between 12 and 31 retinae per group (see Supplementary information for full details). *BF*_inclusion_ < 0.33 was considered as the evidence for null hypothesis (i.e., the data are at least three times as likely under the null hypothesis than the alternative).

The four molecules were labelled throughout the retina, but no difference was observed between wild-type and *pdzk1*-KO fish at the ages assessed (Figure 5a–d′; Supplementary Figure S3). Similarly, two-way ANOVA of puncta density in IPL did not reveal any differences between groups (Figure 5e–h; Supplementary Table S2). Bayesian ANOVA [28] provided support for the null hypothesis of no difference for serotonin and NMDAR1 puncta densities across all tested ages (*BF*_inclusion_ = 0.18 and *BF*_inclusion_ = 0.26, respectively; Figure 5f–g; Supplementary Table S2), suggesting these two molecules are particularly unlikely to underlie the altered retinal function.

## 4. Discussion

In this study, 7-dpf *pdzk1*-KO fish showed spatial-frequency tuning functions of higher amplitude compared to wild-type (Figure 1). The human *PDZK1* gene is located on chromosome 1q21.1, a high-risk region for schizophrenia [29] and autism spectrum disorders (ASD) [30]. It has been linked to altered contrast sensitivity, a visual endophenotype of both neurodevelopmental disorders, with each additional copy of the minor allele of the rs1797052 polymorphism in the regulatory region of *PDZK1* increasing sensitivity by more than half a standard deviation [3]. Our data for enhanced contrast sensitivity in *pdzk1*-KO larvae are consistent with these observations and highlight the cross-species conservation of gene function.

Our measurement of contrast-response functions suggests that the improved spatial-frequency tuning function in *pdzk1*-KO mutants is likely due to abnormal gain control (Figure 2). Changes in gain control can manifest in the contrast-response function in three ways. First, *baseline control* vertically shifts the entire function without changing the function shape, indicating contrast-independent modulation of the response. Second, *contrast-gain control* horizontally shifts the entire function without changing the function shape, indicating changes of contrast threshold for eliciting any response (i.e., contrast sensitivity). Finally, *response-gain control* is characterised by changes in the slope or maximum response without changing the baseline of the function [31, 32]. In our data, responses were not saturated even at full contrast, making the slope of the function the best reflection of response gain. However, we found no evidence of response gain changes in knockout fish. Instead, mutants showed significantly lower contrast thresholds (at 25 and 50°/s) compared to wild-type fish (Figure 2e; Supplementary Table S4), indicating abnormal contrast-gain control in mutants. This is consistent with the association between the human 1q21.1 genomic region containing *PDZK1* and risk of autism and schizophrenia: Like our *pdzk1*-KO larvae, infants at high-risk for autism (i.e., those who have an older sibling diagnosed with autism) and people at high-risk for psychosis (i.e., those who fulfil the criteria of Attenuated Psychotic Symptoms and Brief Limited Intermittent psychotic Symptoms [33]) also show higher luminance contrast sensitivity [34, 35]. Altered contrast gain may originate in the retina via reduced inhibitory signals from amacrine cells onto parasol ganglion cells [36], or it may occur in higher visual regions; however, the absence of a visual cortex in zebrafish, along with the electrophysiological findings of the present study, suggests the effects of *PDZK1* variation are more likely retinal in origin.

Our ERG results further corroborate that the neurodevelopmental risk gene *PDZK1* contributes to retinal function. With enhanced spatial contrast-sensitivity in *pdzk1-KO* larvae compared to wild-types, one might expect *larger* ERG responses for mutants, but we found a reduction in the *b*-wave. This reduction accords with the reduced scotopic ERG *b*-wave reported in individuals with schizophrenia and ASD [37, 38]. Individuals with schizophrenia exhibit significantly decreased *a*-wave amplitudes [37]; however, the *b*-wave reduction in *pdzk1*-KO zebrafish was not due to photoreceptor deficits, as *a*-wave amplitudes did not differ between *pdzk1*-KO and wild-type larvae across all flash intensities (Figure 3b, Supplementary Table S5). In contrast, the ERG phenotype observed in *pdzk1*-KO mutants may be more similar to those with ASD, who show no significant changes in *a*-wave amplitude [38], although further research into the developmental trajectory of the ERG changes is needed.

Selective attenuation of the *b*-wave at higher stimulus intensities (1.89 and 2.48 log cd.s.m^−2^) in *pdzk1*-KO fish suggests that *pdzk1* disruption affects either bipolar cells directly or via third-order neurons (i.e. amacrine and ganglion cells). If enhanced behavioural contrast sensitivity in *pdzk1*-KO zebrafish arises from deficits of inhibitory mechanisms, altered bipolar cell responses underlying the *b*-wave may be modulated by a more generalised change in retinal inhibition. In rodents, antagonist blockage or genetic knockout of the GABA_c_ receptor, a key inhibitory receptor on bipolar cells, leads to a reduced ERG *b*-wave but increased spontaneous and light-evoked activity in retinal ganglion cells [39-41]. How PDZK1 might be involved in the crosstalk between bipolar, amacrine and ganglion cells remains to be investigated.

While the observed *b*-wave deficit points to bipolar or third-order retinal dysfunction, there could be other potential causes. PDZK1 may contribute to microvillus formation in the retinal pigment epithelium (RPE) through its interaction with ezrin/radixin/moesin (ERM), binding phosphoprotein 50 kDa (EBP50) and ezrin [42, 43]. RPE apical microvilli interdigitate with outer segments of adjacent photoreceptors, involved in essential functions of the RPE for retinal homeostasis (e.g., phagocytosis of photoreceptor debris, nutrient transportation into retina, removal of photoreceptor waste products and visual pigment regeneration) [42, 44]. Therefore, loss of PDZK1 may disrupt retinal homeostasis and thus retinal function; however, in this case, one might also expect altered *a*-wave responses, which were not observed. Knockout of murine *Pdzk1* leads to reduced post-transcriptional expression of scavenger receptor class B type I (SR-BI), altering lipid metabolism for high-density lipoprotein (HDL) in total plasma [45]. Plasma HDL has been associated with age-related macular degeneration [46, 47], linking PDZK1 via SR-BI to a pathogenic mechanism of eye disease. PDZK1 may also maintain neural homeostasis by increasing the activity of glutamate transporter excitatory amino acid carrier (EAAC1) on the cell membrane [48]. Cross-talk between EAAC1, NMDARs and α-amino-3-hydroxy-5-methyl-4-isoxazolepropionic acid (AMPA) receptors tightly modulates the extracellular concentration of glutamate to avoid excitotoxicity [49]. In mice, EAAC1 deficiency results in excitotoxic retinal damage, specifically ganglion cell loss [50]. Loss of PDZK1 might thus lead to dysfunction of the inner retina by this indirect route.

Our histological analyses indicated that general retinal morphology (IPL thickness, retinal size and retinal thickness) and the density of our four molecules of interest (PSD-95, serotonin, NMDAR1 and CRFR1) in the IPL remain unchanged in mutant retinae (Figure 5f–g; Supplementary Table S2). We were not able to examine an effect of *pdzk1* knockout on receptors in the OPL, as the zebrafish OPL is too thin for effective quantification with the available imaging approaches. Previous studies have shown that early in disease progression (e.g., in diabetes) or under physical stress (e.g., high intraocular pressure), ERG abnormalities can manifest without apparent structural changes [51, 52]. However, we analysed molecular density from 7 dpf to 8 wpf, suggesting that progressive changes in morphology or the tested molecules are unlikely to underlie altered function in *pdzk1*-KO mutants.

We have confirmed the absence of a genetic mechanism for histological-phenotype rescue for *pdzk1*-KO mutants, so we are confident these histological results are robust. Previous studies have reported that alternative splicing of the mutated mRNA frequently occurs in CRISPR-generated mutants by skipping the whole exon of the target locus and may produce functional or partially functional proteins [53, 54]. However, in our case, variant splices of *pdzk1* mRNA are absent in mutant fish, with a single *pdzk1* mRNA expressed with the stop-codon cassette, leading to a premature, non-functional Pdzk1 (Supplementary Figure S4b–e). Recent studies have also raised concerns that mutant mRNA degradation due to nonsense-mediated decay (NMD) can activate compensatory genetic mechanisms, leading to transcriptional adaptation in genetically modified models [55]. Multi-level analysis (OMR, ERG and histology) of *pdzk1*-KO mutants treated with NMD interference (NMDi14; see Supplementary methods for full information) demonstrated that genetic compensation does not occur in our *pdzk1*-KO fish (Supplementary Figure S5 and Table S7).

Ultimately, our histological data provide evidence that while PDZK1 may associate with synaptic proteins, it is not necessary for their clustering into synaptic puncta. Thus, the molecular mechanisms underlying the contribution of PDZK1 to synaptic circuits and visual processing remain unclear; further research may focus on other PDZK1-interacting molecules such as Synaptic Ras GTPase-activating protein 1 (SynGAP1) and Kelch-like protein 17 (KLHL17). As we only surveyed gross retinal morphology, higher-power imaging of individual subtypes of neurons may reveal very specific contributions of PDZK1 to the maintenance of cellular morphology and synaptic distribution within select visual pathways.

## 5. Conclusion

Our study corroborates the association between *PDZK1* and visual function. Using OMR, we showed that *pdzk1*-KO mutants had increased amplitude of spatial-frequency tuning functions, likely due to altered contrast gain control. Knockout zebrafish also exhibited reduced ERG *b*-wave amplitude, indicating deficits in the inner retina (i.e., bipolar, amacrine and ganglion cells). We found no morphological changes nor changes in density of four PDZK1-interacting molecules in *pdzk1*-KO mutants. *Pdzk1*-KO mutants showed similarities in visual phenotype to individuals with, or at high-risk for, neurodevelopmental disorders, especially ASD [35, 38]. Using a range of analytical approaches and targeted gene editing, the zebrafish model provides a platform for assessing the role of genes associated with abnormal visual function.

## Acknowledgements

The authors wish to thank the staff of the zebrafish facility at the Walter and Eliza Hall Institute of Medical Research for animal maintenance, the Biological Optical Microscopy Platform (BOMP) of the Melbourne Advanced Microscopy Facility for providing instruments for confocal imaging, and Prof. Steven D. Leach of Johns Hopkins University for providing the Tg(*ptf1a*:*GFP*) zebrafish line.

## Contributions

JX contributed to conception and design, carried out data collection, participated in data analysis, drafted and critically revised the manuscript. PRJ contributed to conception and design, supervised histological data collection and generation of mutants, and critically revised the manuscript. BVB contributed to conception and design, supervised electrophysiological data collection, and critically revised the manuscript. SD carried out generation and genotyping of mutants and critically revised the manuscript. PTG contributed to conception and design, supervised behavioural data collection, participated in data analysis, and critically revised the manuscript. All authors gave final approval for publication.

## Competing interests

The authors declare no competing interests.

## Funding

This work was supported by a grant from the Melbourne Neuroscience Institute (to PTG, PRJ & BVB). PTG was supported by a Discovery Early Career Researcher Award from the Australian Research Council (DE160100125). JX was supported by a Melbourne Research Scholarship.

## Data availability

The datasets supporting this article are available on the Open Science Framework (https://osf.io/s49z6).

## Notes

https://osf.io/s49z6

### Competing Interest Statement

The authors have declared no competing interest.

