## Supplementary Information for "Altered visual function in a larval zebrafish knockout of neurodevelopmental risk gene *pdzk1*"

Supplementary information for
**Altered visual function in a larval zebrafish knockout of neurodevelopmental risk gene *pdzk1***

Jiaheng Xie, Patricia R. Jusuf, Bang V. Bui, Stefanie Dudczig, Patrick T. Goodbourn*

Supplementary Methods
Supplementary Figures S1–S5
Supplementary Table S1–S7
Supplementary References

Preprint 1.0
22 September 2020

### Supplementary Methods

*Standard RT-PCR*

Recent studies indicate that CRIPSR/Cas9 gene editing can lead to skipping of the whole targeted exon and sometimes exons following, and produce functional or partially functional splice variants of the mutated gene [1, 2]. To confirm that the *pdzk1* gene has no functional splice variants except for that targeted by our CRISPR for insertion of the stop-codon cassette, we used standard RT-PCR. Twenty larvae per genotype were used for RNA extraction using the RNeasy Mini Kit (QIAGEN, Hilden, Germany). Copy DNA was generated by reverse transcription using a Tetro cDNA synthesis kit (Bioline, London, UK). The zebrafish *pdzk1* gene has nine exons. Primers were designed against (1) an RNA region across exons 1–6 containing the insertion site (Primer pair 1; *P1*) and (2) a region starting with the stop-codon cassette insertion in exon 2 and ending in exon 6 (Primer pair 2; *P2*; Supplementary Table S1 and Supplementary Figure S4a). PCR conditions were 98 °C for 3 min (1 cycle), 98 °C for 30 s, 56 °C for 30 s and 72 °C for 1 min (35 cycles), and 72 °C for 3 min. Electrophoresis was performed using 3% TAE agarose gel and products were imaged under UV light.

*Genetic compensation test*

*Pdzk1*-KO embryos were randomly assigned to two groups after collection. From 3 dpf, the two groups were either immersed in 0.02% Dimethyl sulfoxide (DMSO) or 10µM NMDi14 (Sigma-Aldrich, St Louis, MA, USA, cat. number SML1538)/0.02% DMSO dissolved in egg water (60 mg/L sea salt), as control or experimental groups, respectively [3]. NMDi14 is a drug for blocking non-sense-mediated decay of mutant mRNA, which is a trigger for genetic compensation. At 7 dpf, the two groups were assessed by OMR for spatial-frequency tuning function, ERG for retinal physiological responses and histology for related molecular density, with the approaches described in the main text. Statistics for OMR, ERG and histological analysis were performed using *F*-tests (a custom MATLAB algorithm), two-way ANOVA with Bonferroni correction and unpaired *t*-test (Prism 8, GraphPad, San Diego, CA, USA; $\alpha = 0.05$), respectively. Sample size for each experiment is detailed in Supplementary Table S7.

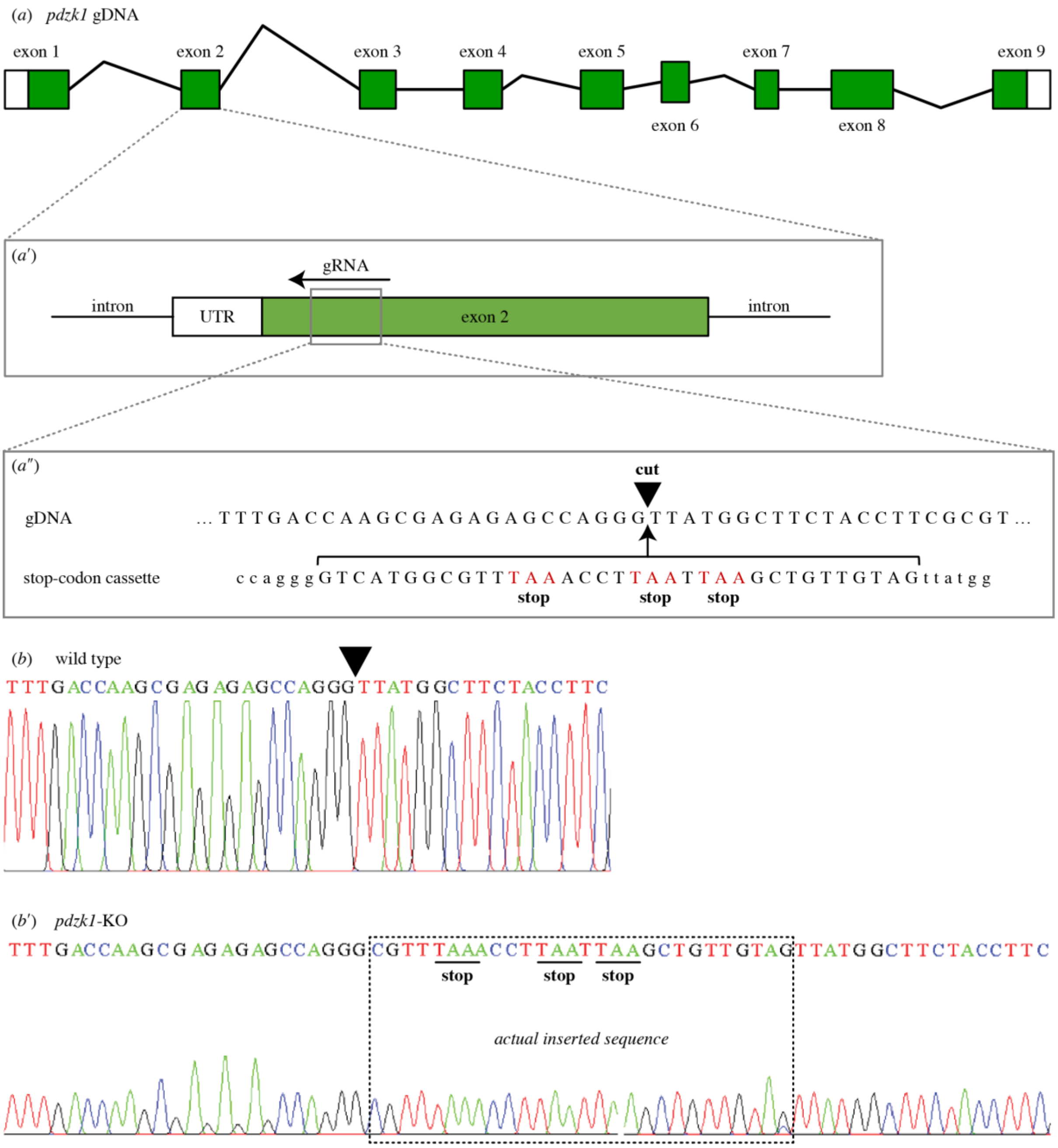


**Supplementary Figure S1**. CRISPR editing of *pdzk1*. (*a*) There are nine exons in *pdzk1* genomic DNA. (*a′*) The guide RNA (gRNA) targets a site in exon 2 (grey box) and recruits the Cas9 enzyme to recognise and cut the DNA. (*a″*) The target cut site (black inverted triangle) has homologous arms matching the stop-codon template that is co-injected with Cas9 and gRNA during the one-cell stage of zebrafish development [4]. In this way, homology-directed repair inserts the stop-codon cassette at exon 2 of *pdzk1*. Stop codons of the injected cassette are highlighted in red. (*b*) Sequence results for wild-type fish. The inverted triangle indicates the targeted cut site. (*b′*) Sequence results for homozygous *pdzk1*-knockout fish. The box shows the sequence inserted by CRISPR gene editing. Sequences on either side were identical to the wild type. Stop codons are highlighted by underline.

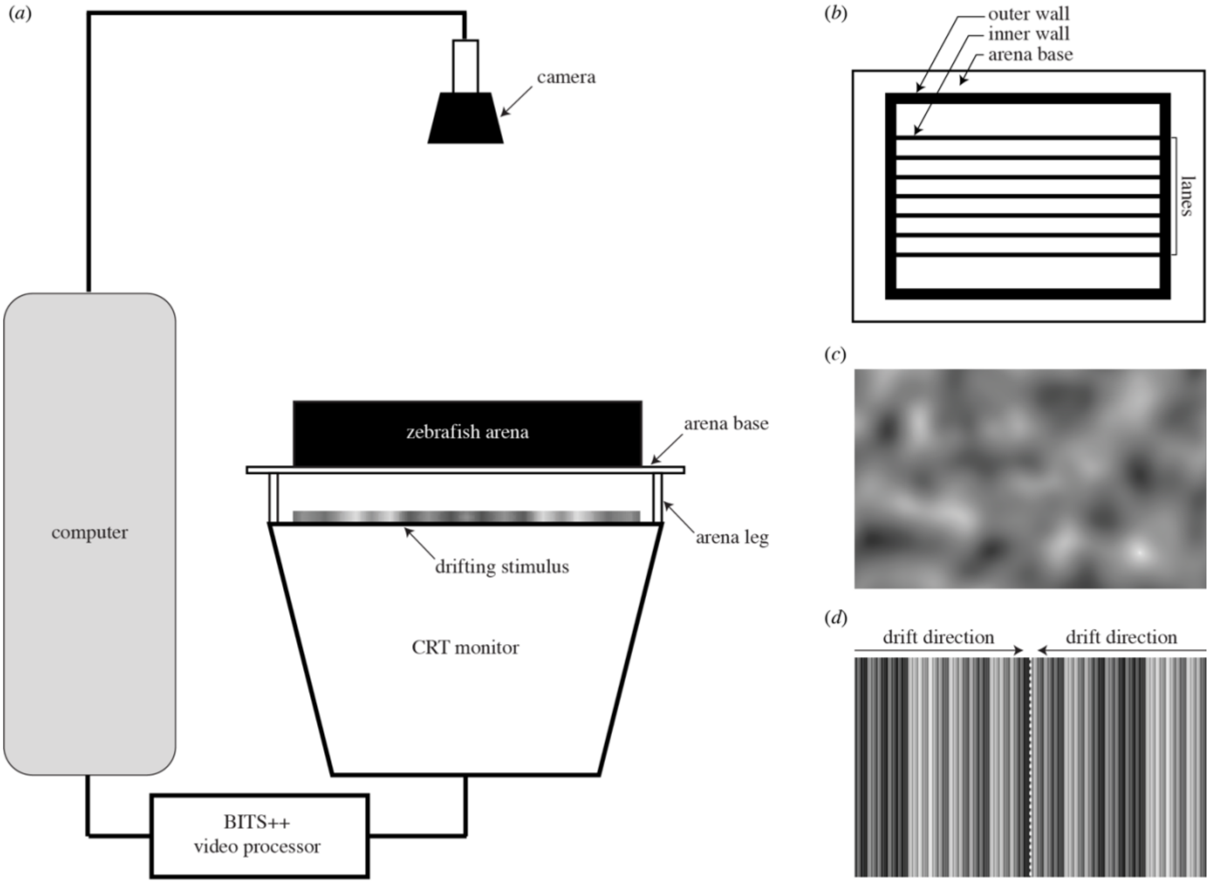


**Supplementary Figure S2**. Schematic of the optomotor assay. (*a*) Optomotor apparatus. Visual stimuli were generated by a Power Mac G5 computer (Apple Inc., Cupertino, CA, USA) running MATLAB R2016b (MathWorks, Natick, MA, USA) with Psychtoolbox extensions [5] and were processed on an ATI Radeon HD 5770 graphics card (AMD, Santa Clara, CA, USA). The outputs were sent to a BITS++ video processor (Cambridge Research Systems, Rochester, UK) for increased contrast resolution, and displayed on a cathode ray tube (CRT) monitor (Model M992, Dell Inc., Round Rock, TX, USA) with its screen facing upwards. Zebrafish were contained in a six-lane arena with a transparent base 62.5 mm above the screen. A C922 Pro Stream webcam (Logitech Company, Lausanne, Switzerland) was fixed 366.5 mm above the base of the arena, controlled by the MATLAB program to take digital images (1080 × 1920 pixels) before and after each trial for calculation of swimming distance. (*b*) Aerial view of the zebrafish arena (290 mm × 25 mm per lane, 50 mm and 10 mm for the outer and inner wall heights, respectively). The inner walls form six independent lanes. (*c*) Example test stimulus, a filtered Gaussian-noise texture. (*d*) Between trials, fish were directed to the centre using a corralling stimulus, in which compound gratings drift from each end, converging on the central line (dashed white line). Figure adapted from [6].

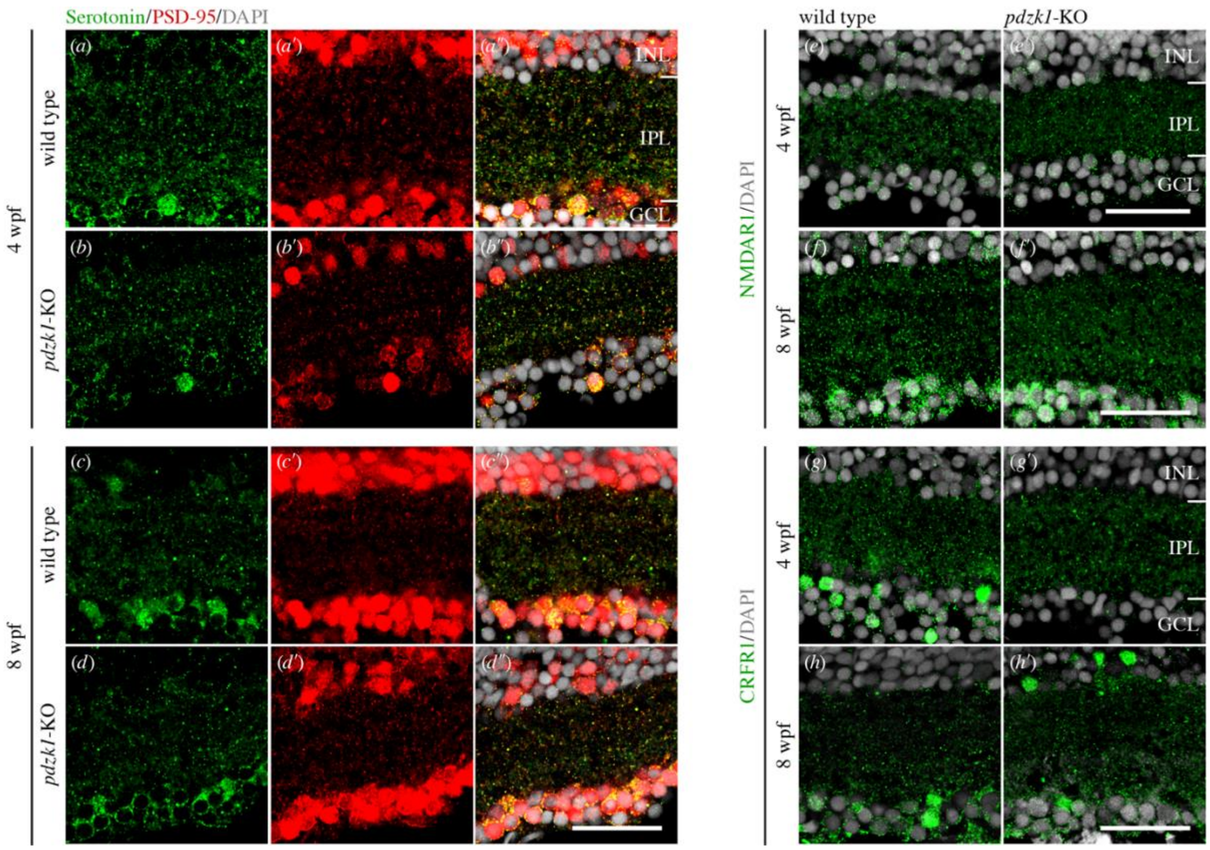


**Supplementary Figure S3.** Micrographs showing immunostaining of PSD-95, serotonin, NMDAR1 and CRFR1 in the IPL of wild-type and *pdzk1*-knockout retinae. Serotonin and PSD-95 were co-labelled in green and red, respectively, in (*a*–*b″*) 4 wpf and (*c*–*d″*) 8 wpf retinae. (*e*–*f′*) NMDAR1 and (*g*–*h′*) CRFR1 were labelled in green in 4 and 8 wpf retinae. For all images, nuclei were stained with DAPI, shown in grey. Scale bars represent 25 µm. PSD-95: post-synaptic density 95; NMDAR1: *N*-methyl-D-aspartate receptor 1; CRFR1: corticotropin-releasing factor receptor 1; INL: inner nuclear layer; IPL: inner plexiform layer; GCL: ganglion cell layer.

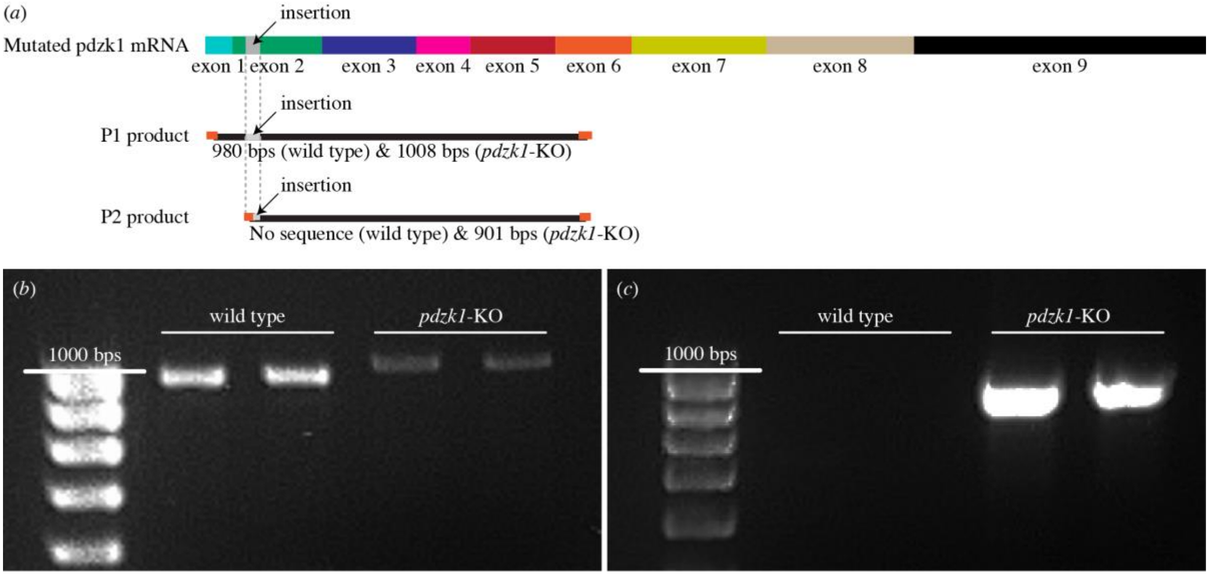


**Supplementary Figure S4**. Standard RT-PCR for testing splice variants (or exon skipping) and expression of *pdzk1* in wild-type and *pdzk1*-knockout (*pdzk1*-KO) mutants. (*a*) Schematics of the two primer pairs for *pdzk1* mRNA and their products. *Pdzk1* mRNA is expressed from the 9 exons of the *pdzk1* genomic DNA. In mutants, the insertion (highlighted in grey on the *pdzk1* mRNA sequence and primer products) should also be expressed within the exon 2 region (green) of the mRNA. Primer pair 1 (P1) was designed to amplify the mRNA region across exons 1–6 containing the insertion, producing 980-base-pair (bp) products for wild type and longer products (1008 bps) for mutants owing to the insertion. Primer pair 2 (P2) was designed to amplify the mutated *pdzk1* mRNA sequence, which starts within the insertion in exon 2 and ends in exon 6. P2 produces 901-bp products only for mutants, with no product for wild-type controls. Orange ends of products represent primers. (*b*) As expected, standard RT-PCR using P1 produced slightly longer products for mutants than for wild type. (*c*) RT-PCR using P2 produced ~900-bp products for mutants but no amplified sequence for wild-type fish. These results demonstrated the expression of the stop-codon cassette in mutants. No splice variants or exon skipping were observed from RT-PCR results.

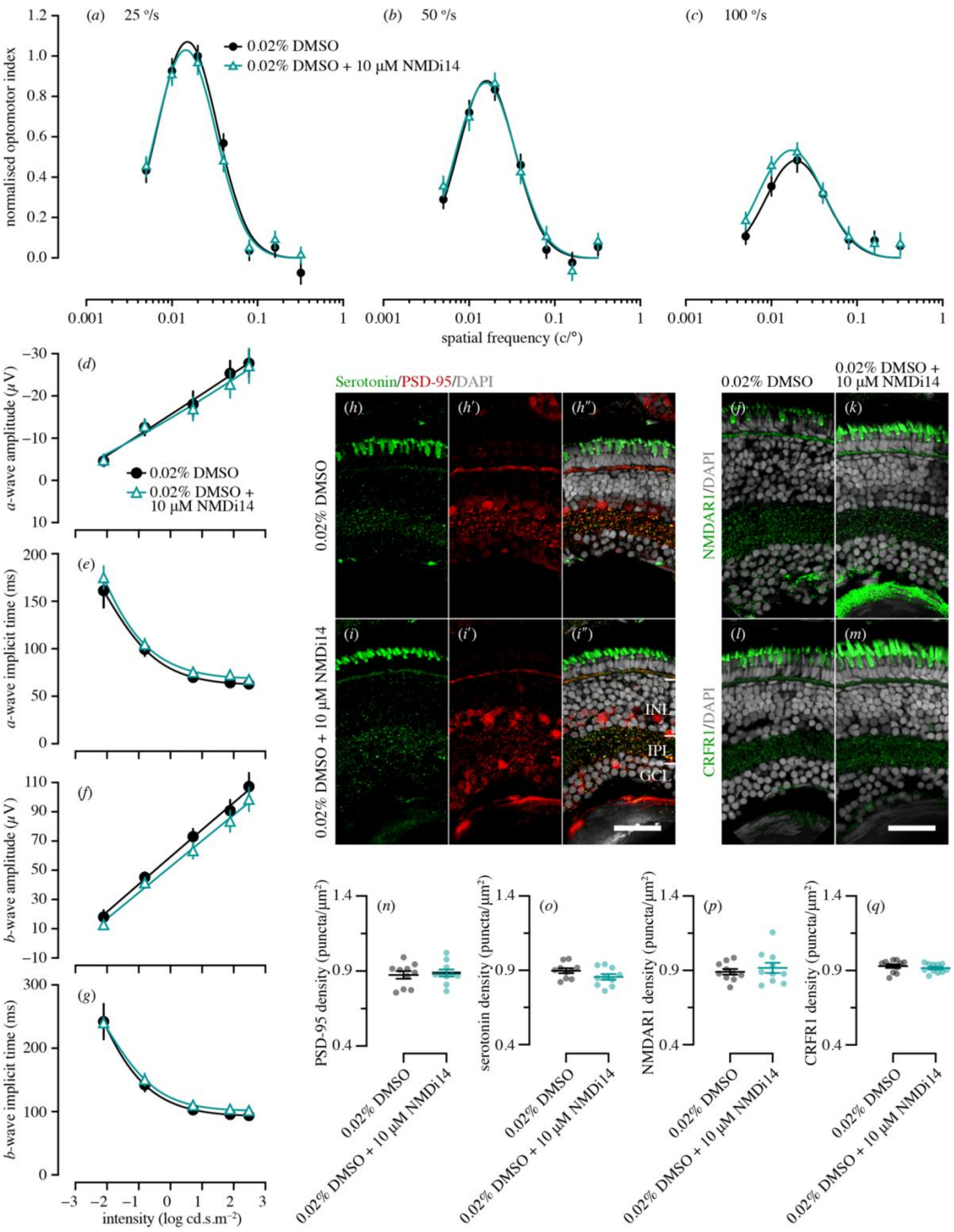


**Supplementary Figure S5.** *Caption overleaf→*

Genetic compensation tests for *pdzk1*-knockout (*pdzk1-KO)* mutants at 7 dpf. Spatial-frequency tunning functions were fitted by a log-Gaussian model for control (*pdzk1*-KO mutants treated with 0.02% DMSO; black circles and lines) and experimental groups (*pdzk1*-KO mutants treated with 0.02% DMSO and 10µM NMDi14; blue triangles and lines) at (*a*) 25, (*b*) 50 and (*c*) 100 °/s. ERG results were shown as the group average (± SEM) of (*d*) *a*-wave amplitude, (*e*) *a*-wave implicit time, (*f*) *b*-wave amplitude and (*g*) *b*-wave implicit time for *pdzk1*-KO mutants treated with 0.02% DMSO (filled black circles) and mutants treated with 0.02% DMSO and 10µM NMDi14 (open cyan triangles). Data were compared using two-way ANOVA; there were no significant differences. Lines were fit by a four-parameter sigmoidal function. For histological assessment, (*h–i″*) serotonin and PSD-95 were co-labelled in green and red, respectively, for both groups. (*j–k*) NMDAR1and (*l–m*) CRFR1 were labelled in green. For all images, nuclei were stained with DAPI, shown in grey. Scale bars represent 25 µm. PSD-95: post-synaptic density 95; NMDAR1: *N*-methyl-D-aspartate receptor 1; CRFR1: corticotropin-releasing factor receptor 1; INL: inner nuclear layer; IPL: inner plexiform layer; GCL: ganglion cell layer. Densities in the IPL of (*n*) PSD-95, (*o*) Serotonin, (*p*) NMDAR1 and (*q*) CRFR1 for mutants treated with 0.02% DMSO (black circles) and mutants treated with 0.02% DMSO and 10µM NMDi14 (cyan circles) were quantified. Points represent data from individual retinae. Thick black lines show group means, and error bars show ± SEM. Unpaired *t*-tests were performed with 10–11 retinae per group; there were no significant differences (see Supplementary Table S7 for full information).

**Supplementary Table S1.** List of oligonucleotide primers.

| Primer pairs | Forward (5′–3′) | Reverse (5′–3′) |
|---|---|---|
| Genomic DNA primers | AGTTTCAGTGTGTTTGTGTTGCA | CACCTAAAGTGTGCACTGTGT |
| Primer for sequencing | GCTGGGGAAAATACTTTCAGACA | |
| P1 | CAACAGACACCTCCTTTACAACA | CACCAGGAGGCAGCATTT |
| P2 | TAAACCTTAATTAAGCTGTTGTAGT | CACCAGGAGGCAGCATTT |

**Supplementary Table S2.** Statistical comparison of PSD-95, NMDAR1, CRFR1 and serotonin density in the inner plexiform layer between wild-type (WT) and *pdzk1*-knockout (*pdzk1*-KO) larvae.

| Age | Comparison | Adjusted *P* value | | | | |
|---|---|---|---|---|---|---|
| | | PSD-95 | Serotonin | NMDAR1 | CRFR1 | |
| 7 dpf | WT vs *pdzk1*-KO | 0.055 | 0.87 | 0.42 | >0.99 | |
| 4 wpf | WT vs *pdzk1*-KO | 0.66 | >0.99 | 0.35 | 0.61 | |
| 8 wpf | WT vs *pdzk1*-KO | >0.99 | >0.99 | >0.99 | 0.14 | |
| PSD-95 density | SS | *df* | MS | *F* | *P* | $BF_{inclusion}$ |
| Interaction | 0.02 | 2 | 0.01 | 1.23 | 0.29 | 0.83 |
| Age | 0.46 | 2 | 0.23 | 27.37 | **<0.0001** | $5.8 \times 10^{7}$ |
| Genotype | 0.03 | 1 | 0.03 | 3.26 | 0.07 | 1.45 |
| Residual | 1.05 | 125 | 0.01 | | | |
| Serotonin density | SS | *df* | MS | *F* | *P* | $BF_{inclusion}$ |
| Interaction | 0.01 | 2 | 0.00 | 0.65 | 0.52 | ***0.16*** |
| Age | 0.50 | 2 | 0.25 | 35.05 | **<0.0001** | $2.2 \times 10^{9}$ |
| Genotype | 0.00 | 1 | 0.00 | 0.16 | 0.69 | ***0.18*** |
| Residual | 0.64 | 93 | 0.01 | | | |
| NMDAR1 density | SS | *df* | MS | *F* | *P* | $BF_{inclusion}$ |
| Interaction | 0.04 | 2 | 0.02 | 2.44 | 0.09 | 0.60 |
| Age | 0.68 | 2 | 0.34 | 42.55 | **<0.0001** | $7.9 \times 10^{10}$ |
| Genotype | 0.00 | 1 | 0.00 | 0.09 | 0.77 | ***0.26*** |
| Residual | 0.79 | 99 | 0.01 | | | |
| CRFR1 density | SS | *df* | MS | *F* | *P* | $BF_{inclusion}$ |
| Interaction | 0.01 | 2 | 0.00 | 1.10 | 0.34 | 0.77 |
| Age | 0.80 | 2 | 0.40 | 94.85 | **<0.0001** | $3.0 \times 10^{15}$ |
| Genotype | 0.01 | 1 | 0.01 | 3.07 | 0.08 | 0.93 |
| Residual | 0.38 | 91 | 0.00 | | | |
| Molecule | Phenotype | Sample size (*N*) | | | | |
| | | 7 dpf | 4 wpf | 8 wpf | | |
| PSD-95 | WT | 31 | 19 | 18 | | |
| | *pdzk1*-KO | 30 | 16 | 17 | | |
| Serotonin | WT | 16 | 20 | 18 | | |
| | *pdzk1*-KO | 12 | 16 | 17 | | |
| NMDAR1 | WT | 14 | 19 | 17 | | |
| | *pdzk1*-KO | 15 | 20 | 20 | | |
| CRFR1 | WT | 14 | 18 | 17 | | |
| | *pdzk1*-KO | 14 | 17 | 17 | | |

*Note:* Two-way ANOVA with Bonferroni correction and Bayesian ANOVA were performed. PSD-95: post-synaptic density 95; NMDAR1: N-methyl-D-aspartate receptor 1; CRFR1: corticotropin-releasing factor receptor 1; *df*: degrees of freedom; SS: sum of squares; MS: mean squares; $BF_{inclusion}$: inclusion Bayes factor. $P < 0.05$ is highlighted in bold text; $BF_{inclusion} < 0.33$ is highlighted in bold italics.

**Supplementary Table S3.** Statistical comparison of spatial-frequency tuning functions.

| Between-group comparison | 25 °/s | 50 °/s | 100 °/s |
|---|---|---|---|
| Omnibus function | $F(3,8) = 14.73$ | $F(3,8) = 10.39$ | $F(3,8) = 5.38$ |
| | $^{**}P = 0.001$ | $^{**}P = 0.004$ | $^{*}P = 0.01$ |
| Amplitude | $F(1,8) = 32.21$ | $F(1,8) = 22.75$ | $F(1,8) = 9.65$ |
| | $^{***}P = 0.0005$ | $^{**}P = 0.001$ | $^{*}P = 0.01$ |
| Peak frequency | $F(1,8) = 0.01$ | $F(1,8) = 1.97$ | $F(1,8) = 0.17$ |
| | $P = 0.92$ | $P = 0.20$ | $P = 0.69$ |
| Bandwidth | $F(1,8) = 0.27$ | $F(1,8) = 0.86$ | $F(1,8) = 0.00$ |
| | $P = 0.62$ | $P = 0.38$ | $P = 0.95$ |

*Note:* 72 trials per group. $^{*}P < 0.05$; $^{**}P < 0.01$; $^{***}P < 0.001$.

**Supplementary Table S4.** Statistical comparison of contrast-response functions.

| Between group comparison | 25 °/s | 50 °/s | 100 °/s |
|---|---|---|---|
| Omnibus function | $F(4,6) = 17.28$ | $F(4,6) = 8.92$ | $F(4,6) = 6.61$ |
| | ***$P = 0.0006$ | **$P = 0.006$ | *$P = 0.01$ |
| Growth rate | $F(1,6) = 0.55$ | $F(1,6) = 0.94$ | $F(1,6) = 0.21$ |
| | $P = 0.47$ | $P = 0.36$ | $P = 0.66$ |
| Contrast threshold | $F(1,6) = 29.17$ | $F(1,6) = 17.25$ | $F(1,6) = 0.38$ |
| | ***$P = 0.0003$ | **$P = 0.002$ | $P = 0.55$ |

*Note:* 36 trials for wild-type and 24 for *pdzk1*-KO groups. *$P < 0.05$; **$P < 0.01$; ***$P < 0.001$.

**Supplementary Table S5.** Statistical analysis of electroretinography (ERG).

| Intensity ($log\ cd.s/m^2$) | Comparison | Adjusted *P* value | | | |
|---|---|---|---|---|---|
| | | *a*-wave amplitude | *b*-wave amplitude | *a*-wave implicit time | *b*-wave implicit time |
| −2.11 | WT vs *pdzk1*-KO | >0.99 | >0.99 | >0.99 | >0.99 |
| −0.81 | WT vs *pdzk1*-KO | >0.99 | >0.99 | >0.99 | >0.99 |
| 0.72 | WT vs *pdzk1*-KO | >0.99 | 0.13 | >0.99 | >0.99 |
| 1.89 | WT vs *pdzk1*-KO | >0.99 | **0.03** | >0.99 | >0.99 |
| 2.48 | WT vs *pdzk1*-KO | 0.99 | **<0.0001** | >0.99 | >0.99 |
| *a*-wave amplitude | SS | *df* | MS | *F* | *P* |
| Interaction | 299.9 | 4 | 74.98 | 0.52 | 0.72 |
| Intensity | 6319 | 4 | 1580 | 10.99 | **<0.0001** |
| Genotype | 319.9 | 1 | 319.9 | 2.22 | 0.14 |
| Residual | 32193 | 224 | 143.7 | | |
| *b*-wave amplitude | SS | *df* | MS | *F* | *P* |
| Interaction | 8060 | 4 | 2015 | 1.84 | 0.12 |
| Intensity | 382106 | 4 | 95527 | 87.31 | **<0.0001** |
| Genotype | 29116 | 1 | 29116 | 26.61 | **<0.0001** |
| Residual | 245091 | 224 | 1094 | | |
| *a*-wave implicit time | SS | *df* | MS | *F* | *P* |
| Interaction | 206.5 | 4 | 51.63 | 0.06 | 0.99 |
| Intensity | 172505 | 4 | 43126 | 53.44 | **<0.0001** |
| Genotype | 2156 | 1 | 2156 | 2.67 | 0.10 |
| Residual | 180752 | 224 | 806.9 | | |
| *b*-wave implicit time | SS | *df* | MS | *F* | *P* |
| Interaction | 2395 | 4 | 598.8 | 0.63 | 0.64 |
| Intensity | 248689 | 4 | 62172 | 65.66 | <0.0001 |
| Genotype | 546.4 | 1 | 546.4 | 0.57 | 0.45 |
| Residual | 212089 | 224 | 946.8 | | |

*Note:* 22 wild-type and 26 *pdzk1*-KO.Two-way ANOVA with Bonferroni correction was performed. $P < 0.05$ is highlighted in bold.

**Supplementary Table S6.** Statistical analysis of retinal morphology.

| Items | Comparison | *P* |
|---|---|---|
| Normalised IPL thickness | WT vs *pdzk1*-KO | 0.81 |
| Normalised retinal size | WT vs *pdzk1*-KO | 0.44 |
| Normalised retinal thickness | WT vs *pdzk1*-KO | 0.58 |

*Note*: Unpaired *t*-test was performed. There were 12 wild-type and 13 *pdzk1*-KO retinae.

**Supplementary Table S7.** Statistical analysis of genetic compensation tests at 7 dpf.

**Spatial-frequency tuning**
(48 trials per group)

| Between-group comparison | 25 °/s | 50 °/s | 100 °/s |
|---|---|---|---|
| Omnibus function | $F(3,8) = 0.33$ | $F(3,8) = 0.10$ | $F(3,8) = 1.40$ |
| | $P = 0.80$ | $P = 0.96$ | $P = 0.31$ |
| Amplitude | $F(1,8) = 0.32$ | $F(1,8) = 0.02$ | $F(1,8) = 0.85$ |
| | $P = 0.59$ | $P = 0.88$ | $P = 0.38$ |
| Peak frequency | $F(1,8) = 0.50$ | $F(1,8) = 0.08$ | $F(1,8) = 1.52$ |
| | $P = 0.50$ | $P = 0.78$ | $P = 0.25$ |
| Bandwidth | $F(1,8) = 0.00$ | $F(1,8) = 0.22$ | $F(1,8) = 0.24$ |
| | $P > 0.99$ | $P = 0.65$ | $P = 0.64$ |

**Electroretinography**
**(**18 ctrl. and 19 exp. retinae; two-way ANOVA with Bonferroni correction; $P < 0.05$ in bold)

| Intensity (log cd.s/m$^2$) | Comparison | Adjusted *P* value | | | |
|---|---|---|---|---|---|
| | | *a*-wave amplitude | *b*-wave amplitude | *a*-wave implicit time | *b*-wave implicit time |
| −2.11 | Ctrl. vs exp. | >0.99 | >0.99 | >0.99 | >0.99 |
| −0.81 | Ctrl. vs exp. | >0.99 | >0.99 | >0.99 | >0.99 |
| 0.72 | Ctrl. vs exp. | >0.99 | >0.99 | >0.99 | >0.99 |
| 1.89 | Ctrl. vs exp. | >0.99 | >0.99 | >0.99 | >0.99 |
| 2.48 | Ctrl. vs exp. | >0.99 | >0.99 | >0.99 | >0.99 |

| *a*-wave amplitude | SS | *df* | MS | *F* | *P* |
|---|---|---|---|---|---|
| Interaction | 460.1 | 4 | 11.78 | 0.08 | >0.99 |
| Intensity | 9126 | 4 | 2281 | 16.07 | **<0.0001** |
| Group | 22.55 | 1 | 22.55 | 0.16 | 0.69 |
| Residual | 22575 | 159 | 142.0 | | |
| ***b*-wave amplitude** | **SS** | ***df*** | **MS** | ***F*** | ***P*** |
| Interaction | 208.2 | 4 | 52.05 | 0.07 | 0.99 |
| Intensity | 136098 | 4 | 34024 | 47.64 | **<0.0001** |
| Group | 1892 | 1 | 1892 | 2.65 | 0.11 |
| Residual | 113566 | 159 | 714.3 | | |
| ***a*-wave implicit time** | **SS** | ***df*** | **MS** | ***F*** | ***P*** |
| Interaction | 337.0 | 4 | 84.25 | 0.14 | 0.97 |
| Intensity | 176158 | 4 | 44039 | 71.06 | **<0.0001** |
| Group | 2205 | 1 | 2205 | 3.56 | 0.06 |
| Residual | 98538 | 159 | 619.70 | | |
| ***b*-wave implicit time** | **SS** | ***df*** | **MS** | ***F*** | ***P*** |
| Interaction | 416.7 | 4 | 104.2 | 0.09 | 0.99 |
| Intensity | 346958 | 4 | 86740 | 70.97 | **<0.0001** |
| Group | 1407 | 1 | 1407 | 1.15 | 0.29 |
| Residual | 194328 | 159 | 1222 | | |

| **Histology** (Unpaired *t*-test) | | |
|---|---|---|
| Molecule | Comparison | *P* |
| PSD-95 density | Ctrl. (*N* = 10) vs exp. (*N* = 10) | 0.71 |
| Serotonin density | Ctrl. (*N* = 10) vs exp. (*N* = 10) | 0.12 |
| NMDAR1 density | Ctrl. (*N* = 10) vs exp. (*N* = 10) | 0.53 |
| CRFR1 density | Ctrl. (*N* = 11) vs exp. (*N* = 11) | 0.37 |

*Note:* Control (ctrl.) and experimental (exp.) groups were *pdzk1*-KO larvae treated with 0.02% DMSO and *pdzk1*-KO larvae treated with 10µM NMDi14 + 0.02% DMSO, respectively. PSD-95: post-synaptic density 95; NMDAR1: N-methyl-D-aspartate receptor 1; CRFR1: corticotropin-releasing factor receptor 1.
